# Euphorbia yadgirensis sp. nov., (Euphorbiaceae) from Karnataka state, India

**DOI:** 10.1101/2022.11.28.518151

**Authors:** N Sarojini Devi, K Raja Kullayiswamy

## Abstract

*Euphorbia yadgirensis* is a new species from the scrub forest of Royangole village, Yadgiri district of Karnataka, India is described and illustrated here. It belongs to the genus *Euphorbia,* subg. *Chamaesyce,* sect. *Longistylae* and resembles *Euphorbia kadapensis*. However, it differs in its decumbent habit (*vs*. erect), sympodial monochasium (*vs*. dichotomous), laciniate stipules (*vs*. scaly), sutures absent on capsule (*vs*. present).

## Introduction

*Euphorbia* L. includes more than 2000 species (Govaerts et al. 2000) which makes it the most diverse genus of the family Euphorbiaceae Juss. and one of the largest genera within the angiosperms (Horn et al. 2012, Webster 2014). Despite its great variation in morphology, ranging from small annual herbs to large trees, the genus is characterized by a synapomorphy unique among angiosperms. The cyathium (Horn et al. 2012) pseudanthial inflorescence consists of a cup-like involucre with glands along its rim (sometimes with appendages of several types) enclosing a single, central pistillate flower (female floret) surrounded by four or five staminate cymules (male floret) with reduced axes. Both pistillate and staminate flowers are highly reduced to a single pistil and single stamen, respectively (Radcliffe-Smith 2001, Prenner and Rudall 2007, Horn et al. 2012, Webster 2014). This may be further reduced in some species with unisexual cyathia, pistillate or staminate flowers to a pistillode or staminodes (Rizzini 1989). Genus *Euphorbia* plants show all known photosynthetic types C_3_, C_4_ and CAM (Webster et al. 1975) and a C_2_ system that represents an early stage of C_3_ to C_4_ transition (Sage et al. 2011).

The subgenus *Chamaesyce* has the second largest species diversity containing around 600 taxa worldwide (Yang et al. 2012). It is the most species-rich lineage of *Euphorbia* in India with 36 percent of the species belonging to this subgenus among 8 subgenera that consist of 35 taxa under five sections: *Chamaesyce, Elegantes, Hypercifoliae, Longistylae* and *Sclerophyllae* (Binojkumar and Balakrishnan 2010) including the recently described *Euphorbia kadapensis* Sarojinidevi and Venkataraju (2014) that fall under the section *Langistylae*. The section *Longistylae* was erected by Binojkumar and Balakrishnan (2010) to accommodate the endemic therophytic Indian taxa with type *E. longistyla* Boissier (1860). This section was reduced to subsection *Hypericifoliae* by Yang et al. (2012) based on molecular data.

While doing intensive exploration of the genus *Euphorbia* from India, the present taxon was collected from the northern part of Karnataka and is here in described as a new species under genus *Euphorbia* subgen. *Chamaesyce,* sect. *Longistylae.*

## Materials and Methods

Specimens were collected from the scrub forest of Royangole village, Yadgiri district, Karnataka state, treated with 80% ethanol, and herbarium specimens were prepared following the standard method (Santapau 1958, Jain and Rao 1977). Flowers and fruits were used for microscopic observations and imaging and preserved in 70% ethanol for future reference. Microscopic images were taken with an Olympus stereo microscope SZX16, DP74 camera attached. Seeds of the species have been provided to the Dharmavana Nature Ark (DNA) for *ex-situ* conservation.

### Taxonomic treatment

#### *Euphorbia yadgirensis* SND. & RKS. *sp. nov.* (Fig. 1, 2 & 3)

*Euphorbia kadapensis* sensu Raja P. et al., JoTT 12(14):17045. 2020.

**Figure 1.**
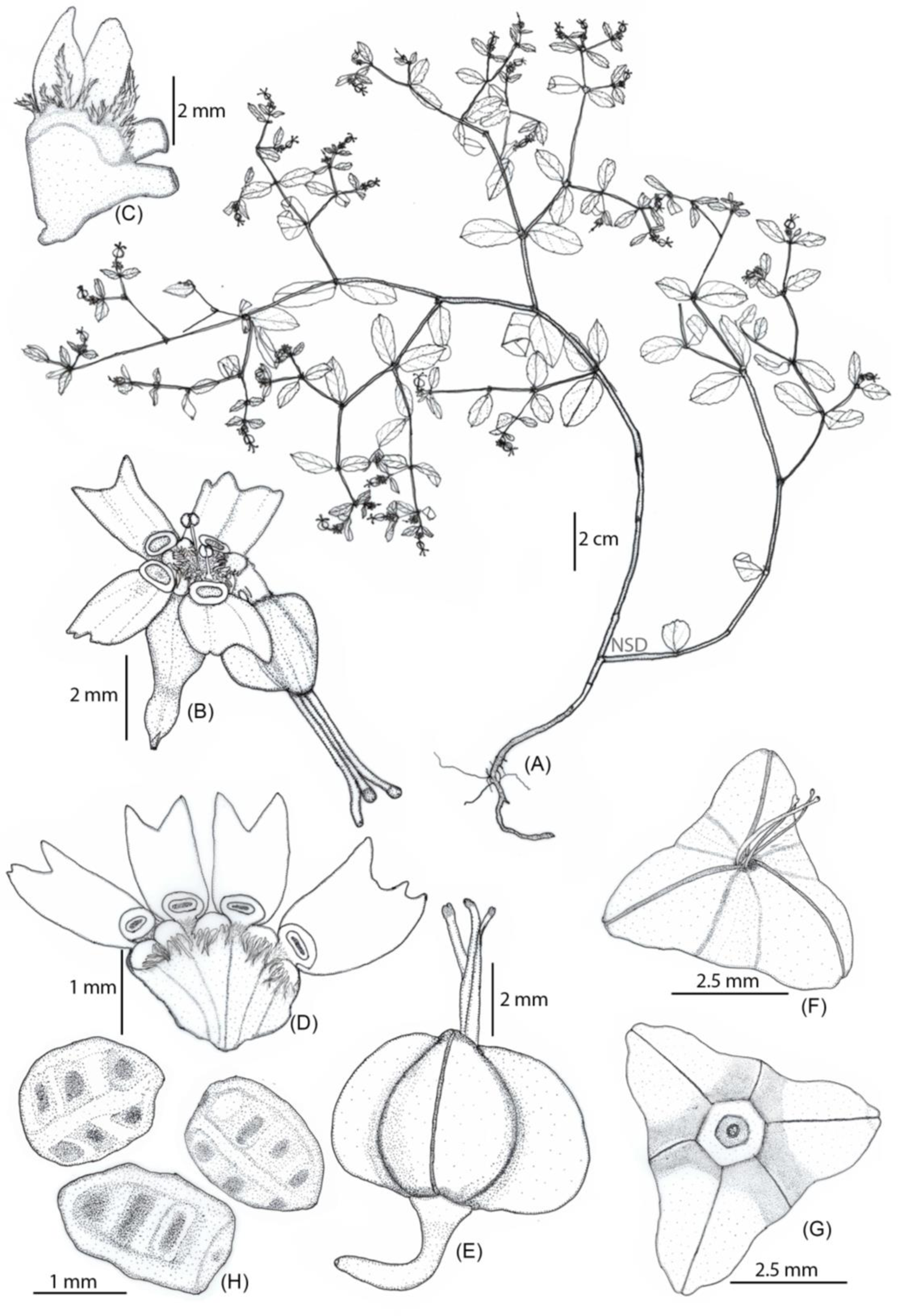
Euphorbia yadgirensis sp. nov. (A) habit, (B) young cyathium, (C) stipules, (D) involucral lobes, limbs and glands, (E) female floret, (F) capsule top view, (G) capsule bottom view, (H) seeds.

**Figure 2.**
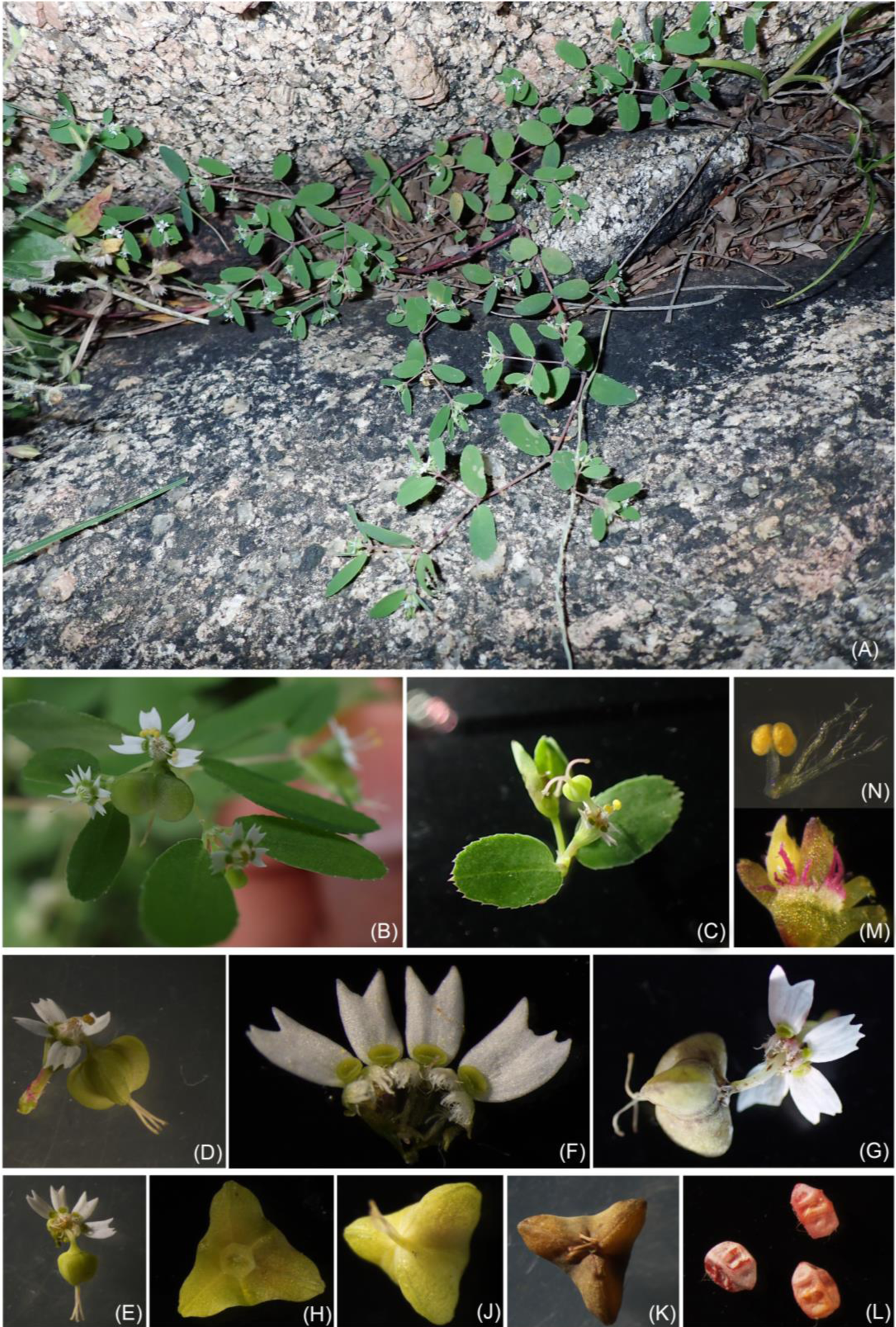
Euphorbia yadgirensis sp. nov. (A) habit, (B) small branch, (C) cyathium at shoot apex, (D) cyathium, (E) cyathium split opened, (F) portion of involucral cup showing lobes, glands and limbs, (G) mature cyathium, (H) capsule bottom view, (J) capsule top view, (K) ripe capsule, (L) seeds, (M) stipule, (N) male floret.

**Figure 3.**
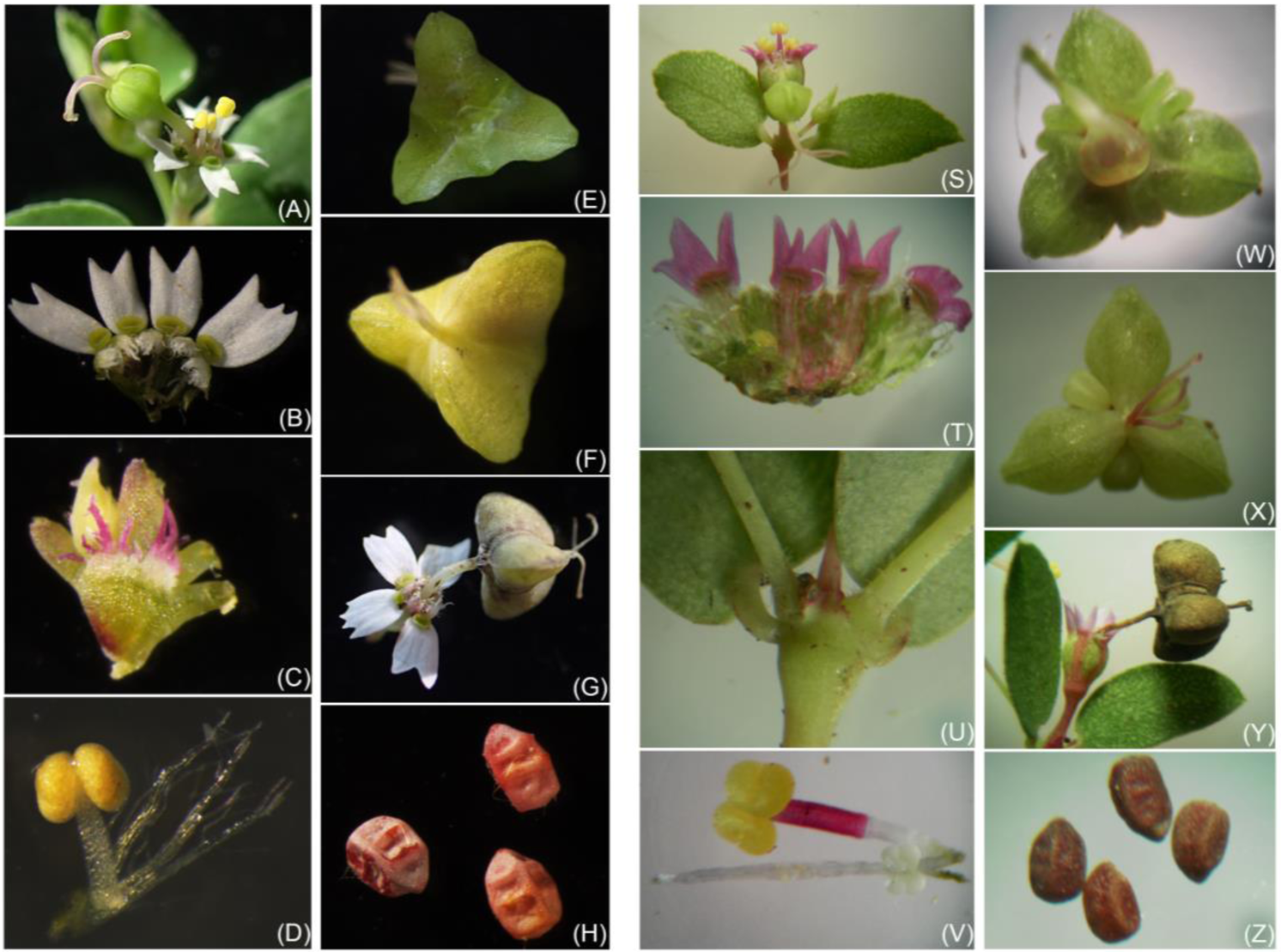
A-H–Euphorbia yadgirensis. (A) cyathium at shoot apex, (B) portion of involucral cup showing lobes, glands and limbs, (C) stipules, (D) male floret (E) capsule bottom view, (F) capsule top view, (G) mature cyathium, (H) seeds. *S-Z–Euphorbia kadapensis.* (S) cyathium at shoot apex, (T) portion of involucral cup showing lobes, glands and limbs, (U) stipules, (V) male floret (W) capsule bottom view, (X) capsule top view, (Y) mature cyathium, (Z) seeds.

##### Type

India (Ind.), Karnataka (KA), Yadgiri district, Shorapur, Royanagole, 16°16’43.36 “ N, 76°24’11.39 “ E, elevation 479 m a.s.l., 11 Oct 2022, RKS HJCB-2090 (holotype MH, iso-BSID, SKU, HDNA).

##### Other specimens

Ind., KA, Royanagole, Yadgiri district, 16.280°N; 76.393°E, alt 484m, 01.ix.2017, P. Raja 2407 at St. Joseph’s college, Tiruchirappalli, Tamil Nadu.

### Diagnosis

*Euphorbia yadgirensis* is an annual decumbent herb growing between rocks in scrub forest, stem glabrous, sympodial-monochasium branching; stipules laciniate, limb of gland bi- or tri-fid one fourth of its length, capsules trigonous prominently keeled, stigmas slightly bifid at apex, seeds transversely deeply rugulose. *Euphorbia yadgirensis* resembles *E. kadapensis* Sarojin. & R.R.V. Raju in filiform branches, elliptic leaves, dissected limb, long style, and trigonous capsule, but differs in habit (decumbent versus erect), height of the plant (20 cm versus 30 cm), branching type (sympodial-monochasium versus dichotomous), stipules (laciniate, 2 mm versus scaly, 1 mm), involucre (3 mm versus 4 mm), limb (bi or trifid one by fourth of its length versus bi or trifid one by two of its length), number of staminate florets (9–12 versus 15–18), bract of male floret (laciniate versus simple), capsules (ridges absent along the sutures versus prominently ridged along the sutures), seeds (transversely deeply rugulose versus transversely slightly rugulose).

### Etymology

The species is named after the type locality of the Yadgiri district in the north of Karnataka state. The type locality is dominated by granite rocks and scrub brush.

### Description

Hemicryptophytic decumbent glabrous herb up to 20 cm tall; latex milky. Stem sympodial-monochasium branching. Branches terete, glabrous, filiform towards apices, internodes 2–3.5 cm long, thickened at nodes; stipules interpetiolar, laciniate, 2 mm long. Leaves simple, opposite, equal, sub-sessile; lamina oblong-elliptic, 2(0.5)–2.5 × 1(0.3)–1.5 cm; base oblong (rounded) [apical leaves characters mentioned in parentheses], margin serrulate, apiculate; mid-nerve prominent, lateral nerves obscure, 4–8 pairs; petiole 3 mm long. Cyathia terminal, up to 10 mm long; peduncles 2.4 mm long; involucre turbinate, 3 mm long, glabrous, green; involucral lobes 5, laciniate, each 5–8-lobed, appendages of glands 4, 2–3-fid, one by fourth of its length, 1.5 mm long, glands 4, oblong, green, stipitate, stalks hairy. Staminate florets 9–12, exserted, 6 mm long, bracteate, bract laciniate, 4 mm long, anther lobes sub-globose, yellow, and dehiscing transversely. Pistillate flowers laterally pendulous, 10 mm long, glabrous, pedicel 4 mm long, ovary tricarpellate, 3 × 5 mm, styles 3, 3.2 mm long, free from the base, stigmas 3, slightly bifid at apex. Capsules schizocarpic, 2.4 mm long, trigonous, distinctly keeled, fruiting pedicels recurved. Seeds 3, ecarunculate, brown, 2 × 1.5 mm, ovoid-oblong, tetragonal, transversely and obscurely furrowed, truncate at base, transversely rugulose.

### Phenology

Flowering and fruiting from September to November.

### Habitat and distribution

*Euphorbia yadgirensis* grows between granite rocks in black gravelly soil, scrub forest near Rayangole, Yadgiri district, elevation ranges from 470 to 480 m a.s.l. The area is dominated by the scrub species *Ziziphus mauritiana* Lam., *Maytenus emarginata* (Willd.) Ding Hou, *Searsia mysorensis* (G.Don) Moffett, and herbs *Tridaxprocumbens* L., *Justiciaprocumbens* L., *Heteropogon contortus* (L.) P.Beauv. ex Roem. & Schult., *Barleriaprionitis* L., *Andrographis longipedunculata* (Sreem.) L. H. Cramer, *Allmania nodflora* (L.) R.Br. ex Wight, *Cyperus rotundus* L. growing along with the present species. Thus far, the species has been collected only from the type locality where there is a threat of grazing and forest fires.

### Comparison

This new species resembles *Euphorbia kadapensis* in its dissected limbs, filiform stem, leaf shape and long style, but differs in many other morphological characters that are tableted below along with another closely related species of this section (table 1 and figure 3).

**Table1:** Morphological comparison of *Euphorbia* sister taxa (the same morphological characters can be seen in Figure 3 and supporting file)

| Characteristics | *Euphorbia yadgirensis* | *E. kadapensis* | *E. longystyla* |
| --- | --- | --- | --- |
| Habit | decumbent | erect | erect |
| Branching | sympodial-monochasium | dichotomous | pseudo-dichotomous |
| Leaf Margin | serrulate from the base | serrulate from the middle | entire or serrate |
| Cyathium | terminal, 10 mm long | terminal and axillary<br>6 – 9 mm long | terminal 8.5 mm long |
| Limb of Gland | 2(3)-fid $\frac{1}{4}$ of its length | 2(3)-fid $\frac{1}{2}$ of its length | Multi-fid from the base |
| Male floret bract | lacinate | simple | simple |
| Stigma | slightly bifid | simple | simple |
| Fruit | 2.4 mm long, without ridges along sutures | 3 mm long, with ridges along sutures | 4 mm long, without ridges along sutures |

### Artificial key to the section *Longistylae* from India

1. Plants hairy, stem ribbed; seeds wrinkled……………… *E. hispida*

– Plants glabrous, stem terete; seeds transversely obscurely furrowed or smooth… (2)
2. Branches always filiform towards the apex; leaves elliptic to oblong-elliptic; styles more than 1.5 mm long……………… (3)

– Branches not filiform towards the apex; leaves oblong or oblong-lanceolate or ovate-oblong; style less than 1.5 mm long……………… (6)
3. Branches minutely velutinous at base; limb of glands dissected……………… (4)

– Branches not velutinous; limb of glands entire, minutely sinuate…… *E. senguptae*
4. Limb of glands deeply multi-fid from the base up to 12-fid……………… *E. longistyla*

– Limb of glands slightly dissected ½ to ¼^th^ of its length, 2–3-fid……………… (5)
5. Plants erect, dichotomously branched; stipules scaly and connate; male floret bract simple; limb of gland bifid or trifid from the middle; capsule with prominent ridges along sutures; stigmas simple……………… *E. kadapensis*

– Plants decumbent, sympodial-monochasium branching; stipules laciniate; male floret bract laciniate; limb of gland bifid or trifid one fourth of its length; capsule without ridges along the sutures; stigmas slightly bifid……………… **E. yadgirensis**
6. Stem stramineous; leaves too larger on stem than branches; limb of glands 1 mm across……………… *E. jodhpurensis*

– Stem pink or pale pink; all leaves are same as main stem; limbs of glands 2 by 1.5 mm……………… (7)
7. Stem unbranched or rarely dichotomously branched; leaves 3 cm or more long; cyathia pubescent……………… *E. katrajensis*

– Stem branched; leaves less than 3 cm long; cyathia glabrous……………… (8)
8. Height of the plant 30–60 cm; leaves silvery white below; limb of glands irregularely and shallowly wavy……………… *E. erythroclada*

– Height of the plant below 30 cm; leaves pale green below; limb of glands entire or slightly wavy…………… (9)
9. Limb of glands white or yellowish; fruits with 2 or 3 prominent wings on cocci……………… *E. notoptera*

– Limb of glands pink, rosy or pinkish white, never with yellow shade; fruits ridged or keeled, but without wings on cocci……………… (10)
10. Leaves oblong to ovate-oblong, sub-acute to obtuse at apex, 5–11 mm long; capsule prominently broadly and obtusely keeled, distinctly ridged along sutures……………… *E. concanensis*

– Leaves oblong-lanceolate, acute to sub-acute at apex, 10–25 mm long; capsule faintly narrowly and acutely keeled, not ridged along the sutures…. *E. deccanensis*

## Supporting information

Supplementary data for readers

## Acknowledgements

The authors are grateful to the Dharmavana Nature Ark for encouragement and providing the necessary facilities and resources. Thanks to Dr. Kanchian Gandi, of Harvard University for the confirmation of the specific epithet. Also our thanks to Dr. Jahnavi Joshi, Senior Scientist, CCMB for permission for microscopic imaging.

## Figure legends

*Euphorbia longistyla* Bioss. images for reviewer’s understanding purpose

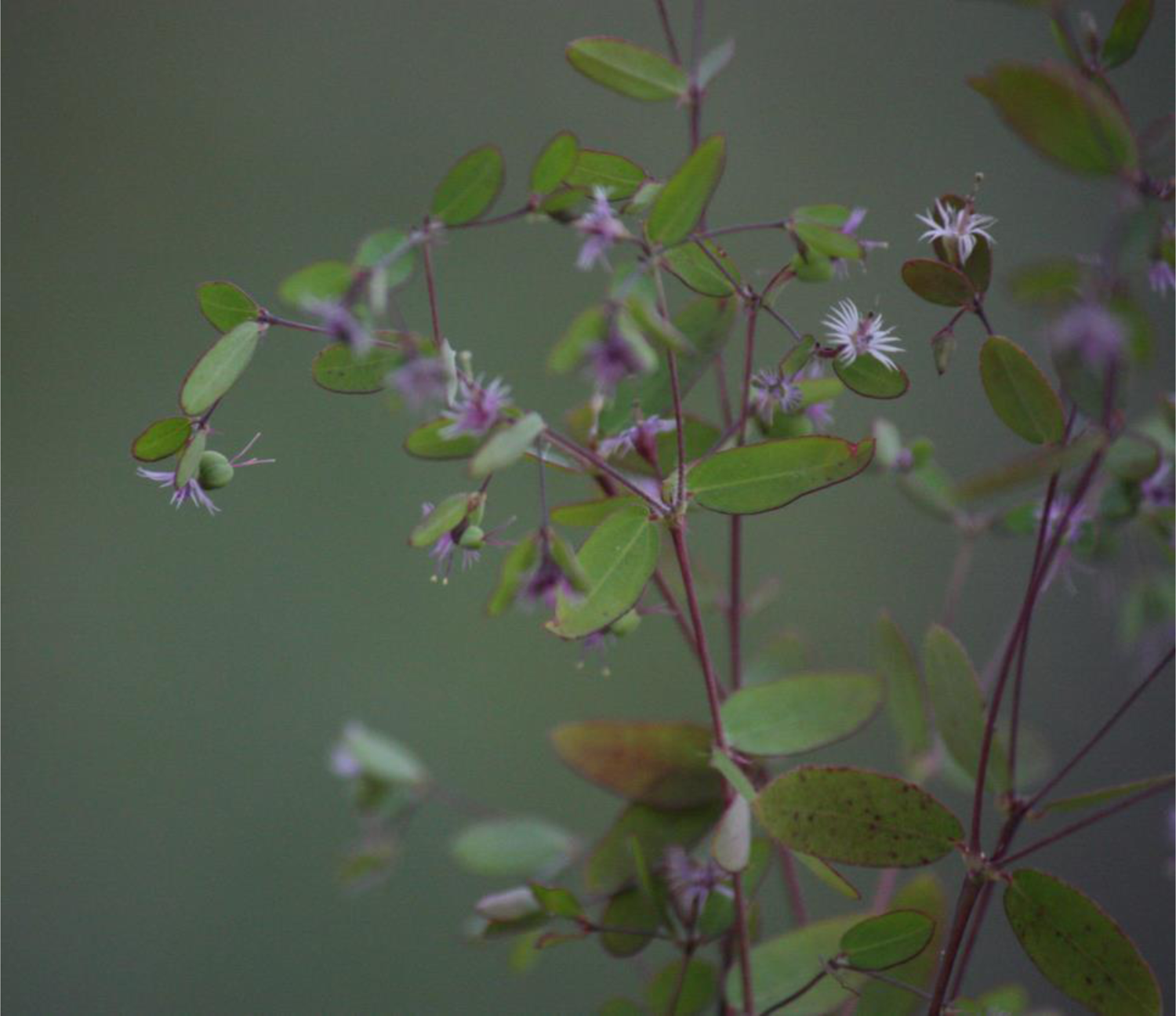
Habit

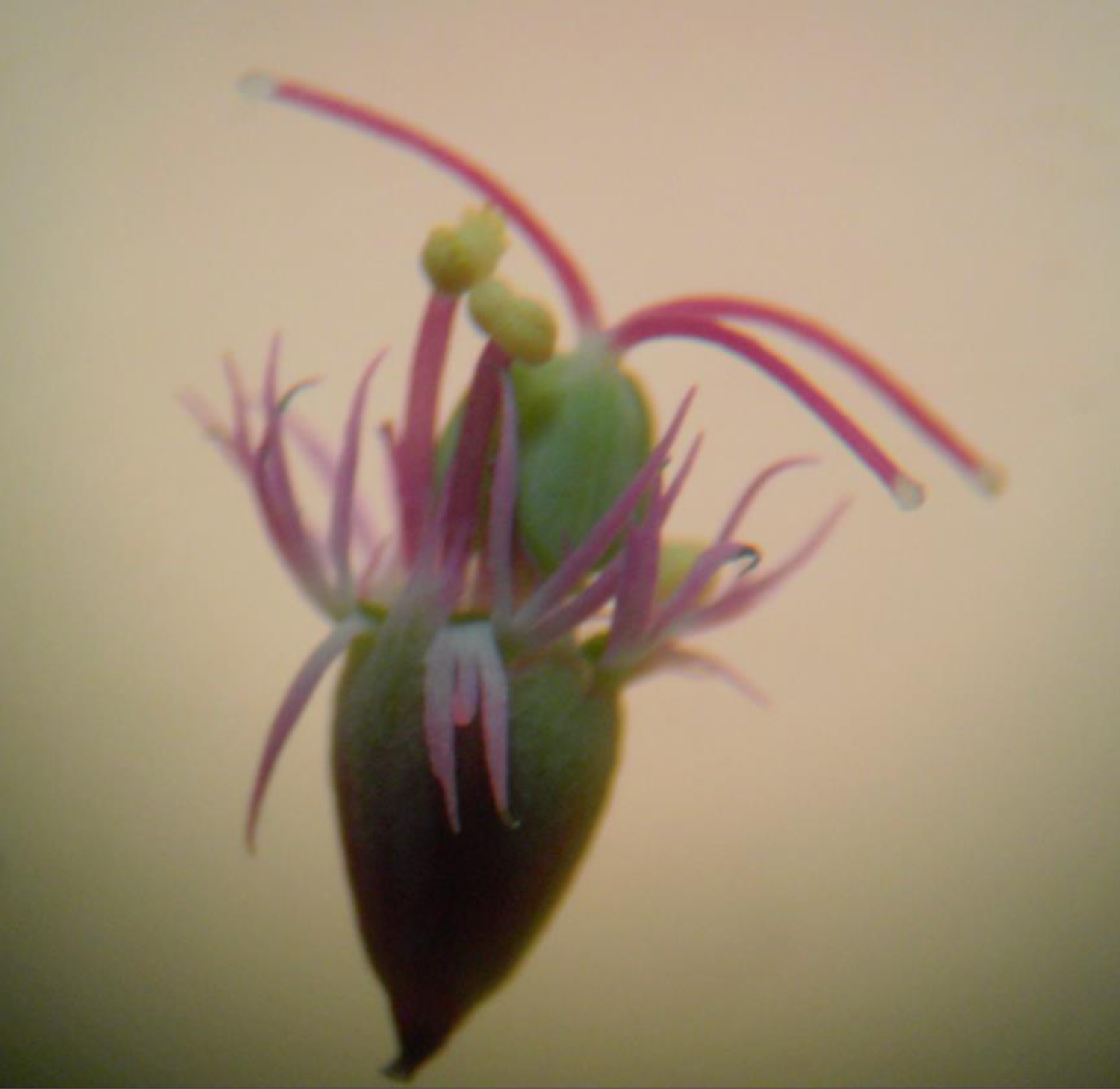
Young cyathium

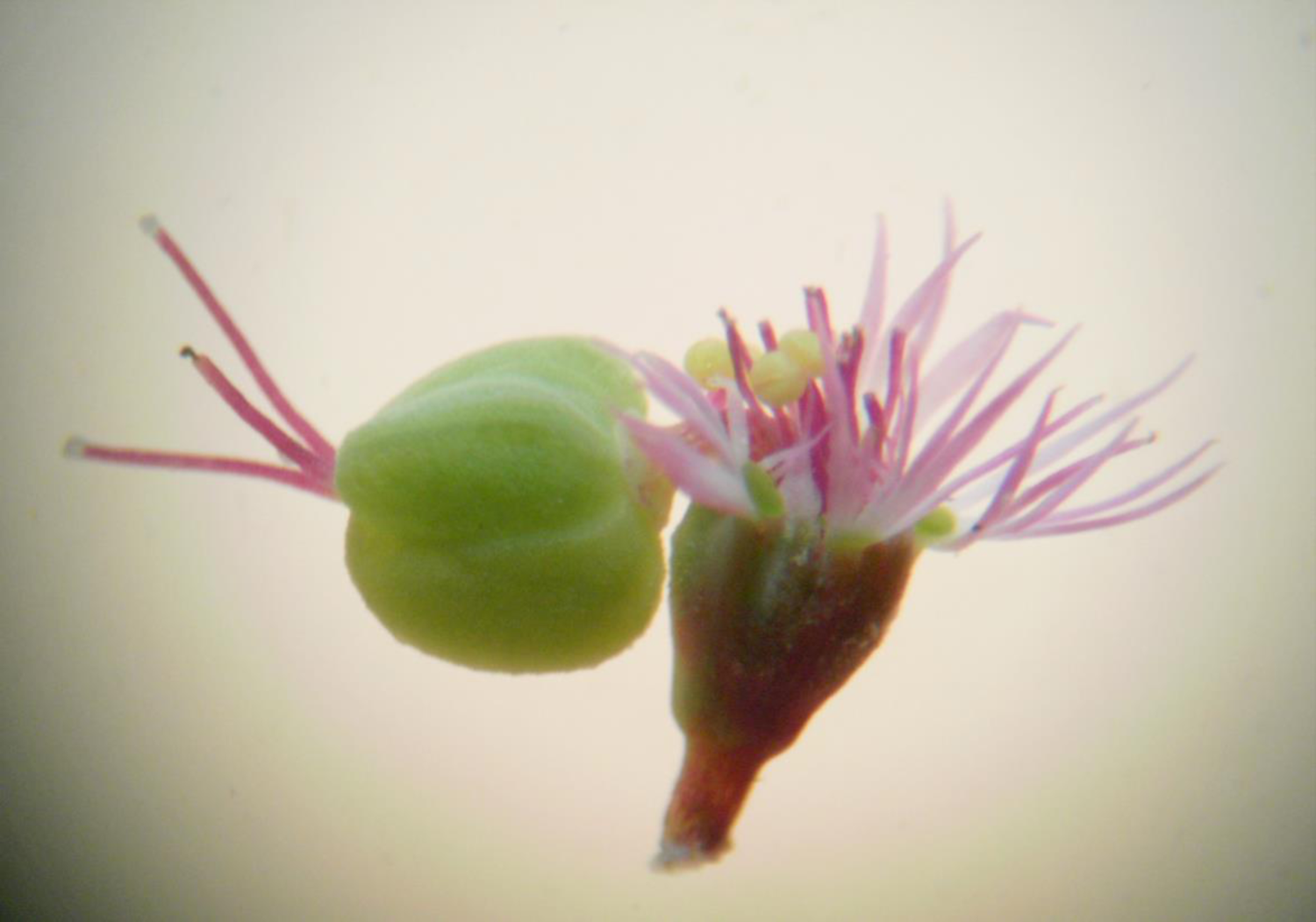
Mature cyathium

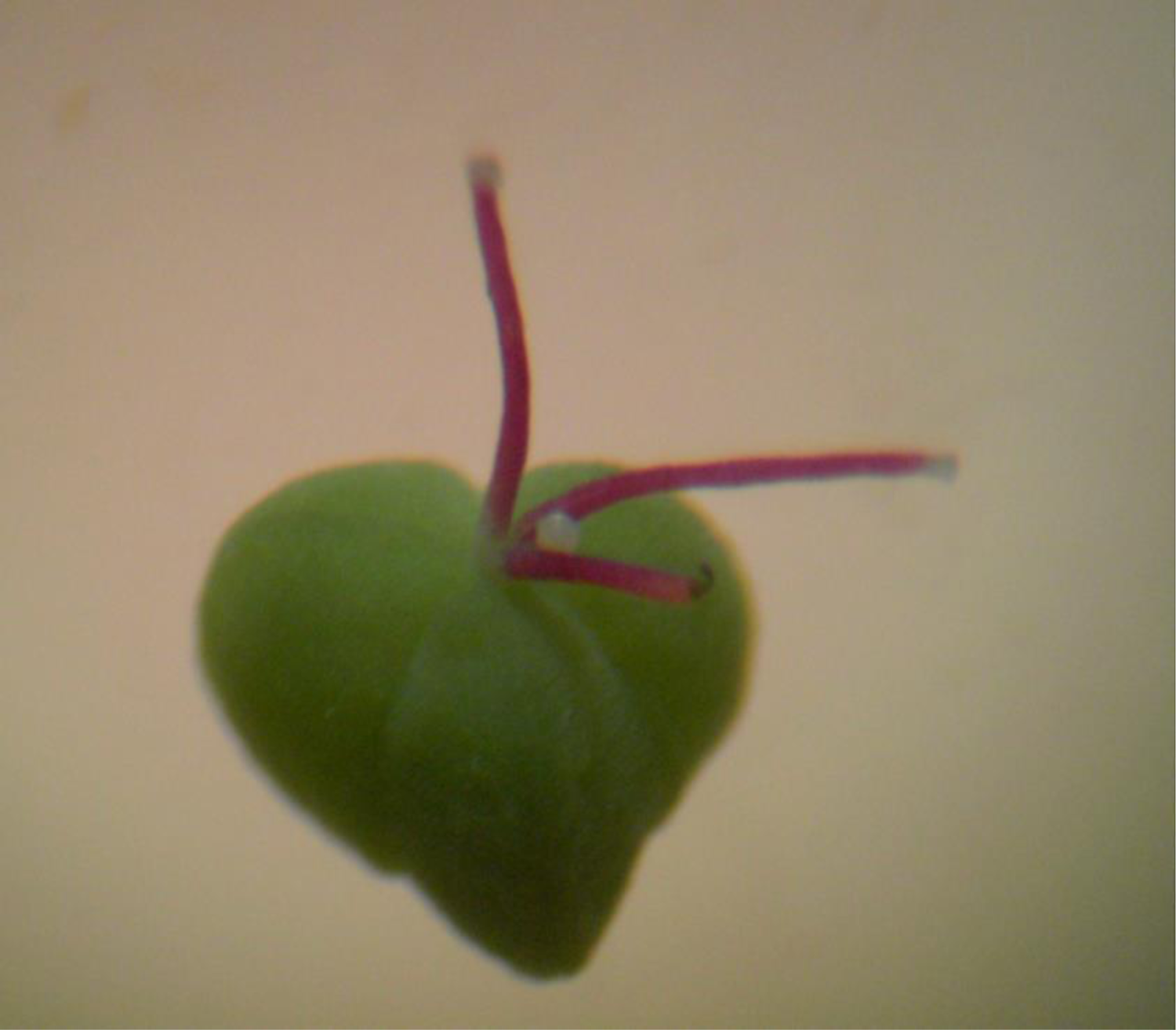
Capsule

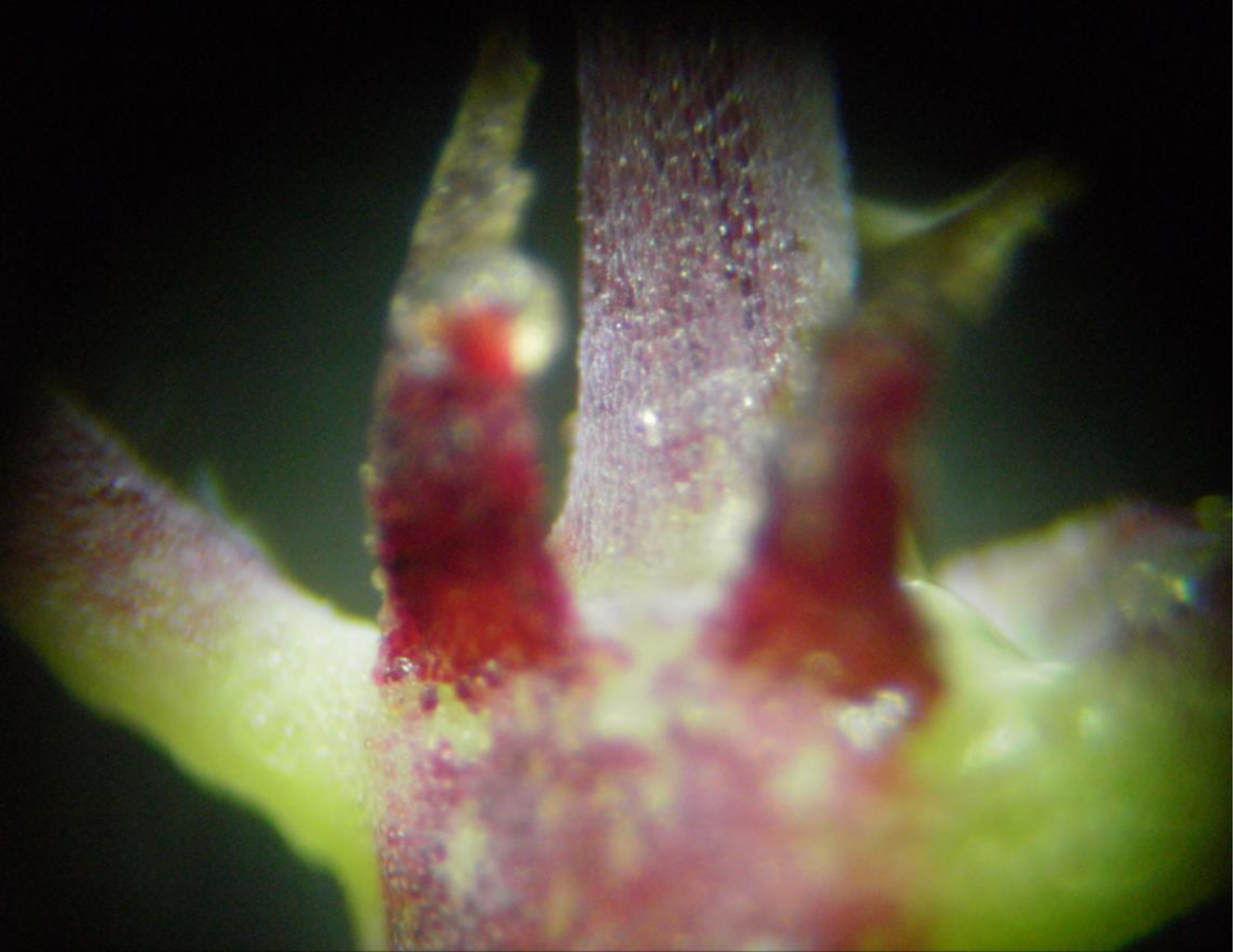
Stipules

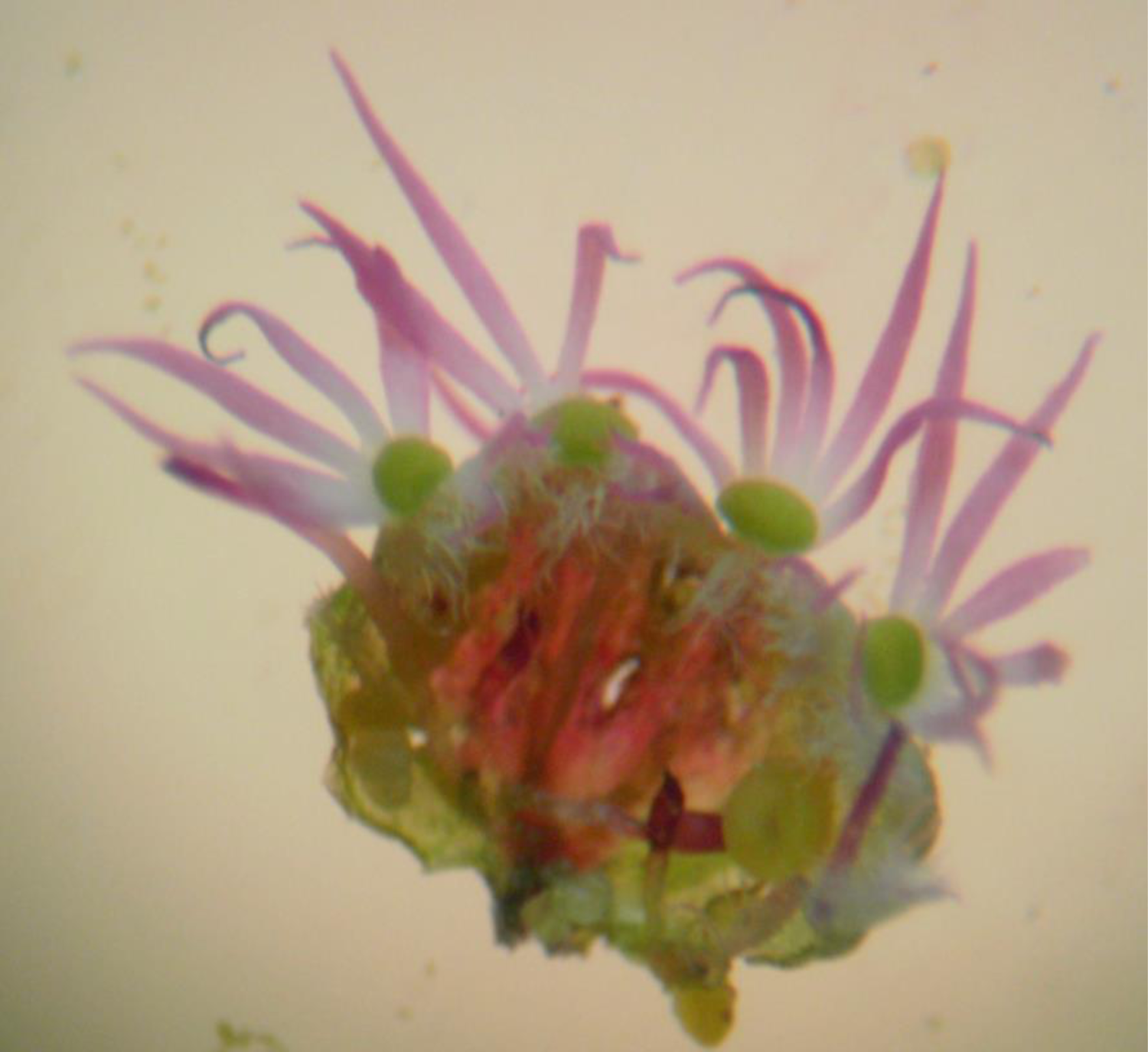
Portion of involucral cup showing lobes, glands and limbs

