## Supplementary data for readers for "Euphorbia yadgirensis sp. nov., (Euphorbiaceae) from Karnataka state, India"

*Euphorbia longistyla* Bioss. images for reviewer’s understanding purpose

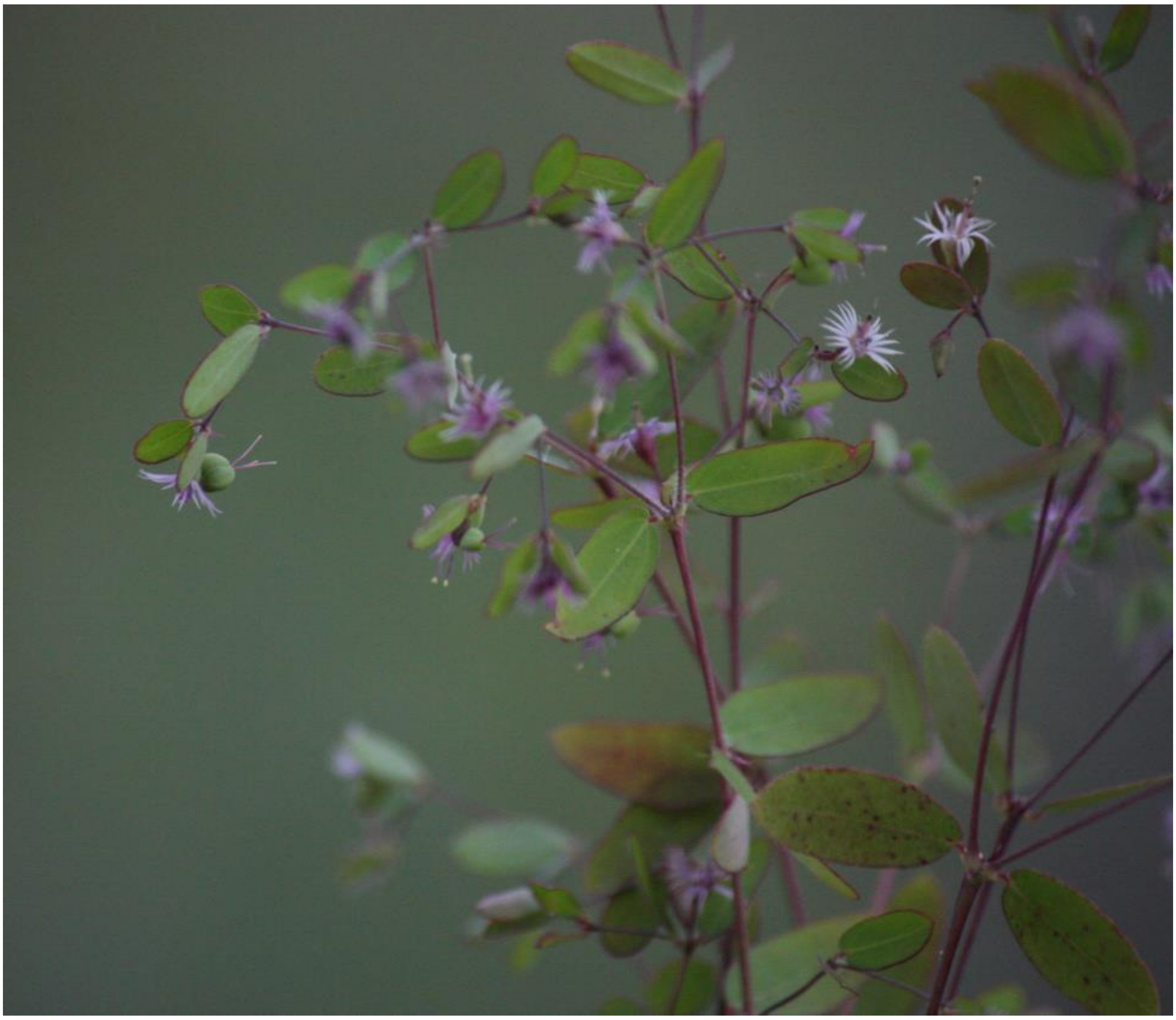

Habit

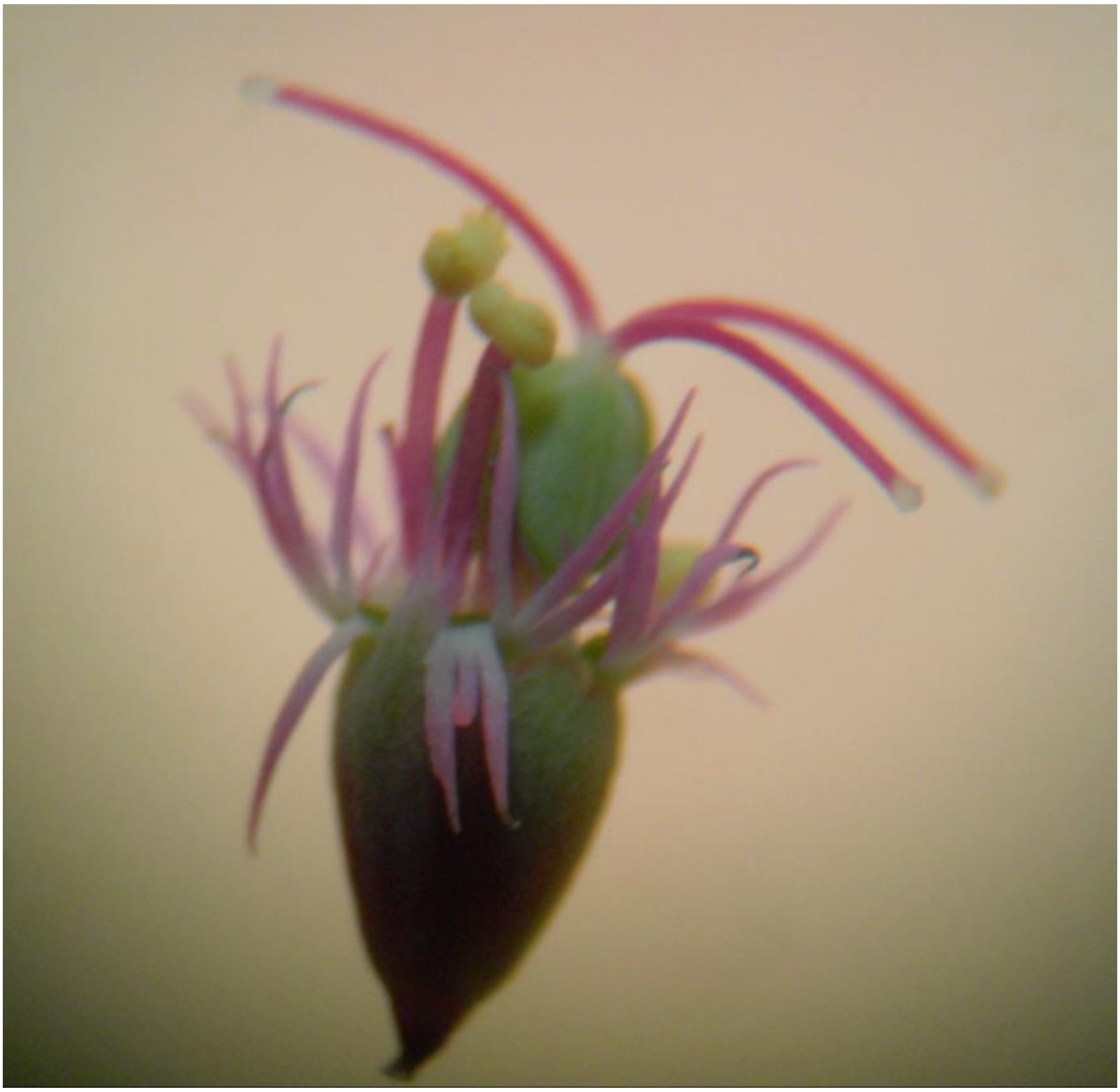

Young cyathium

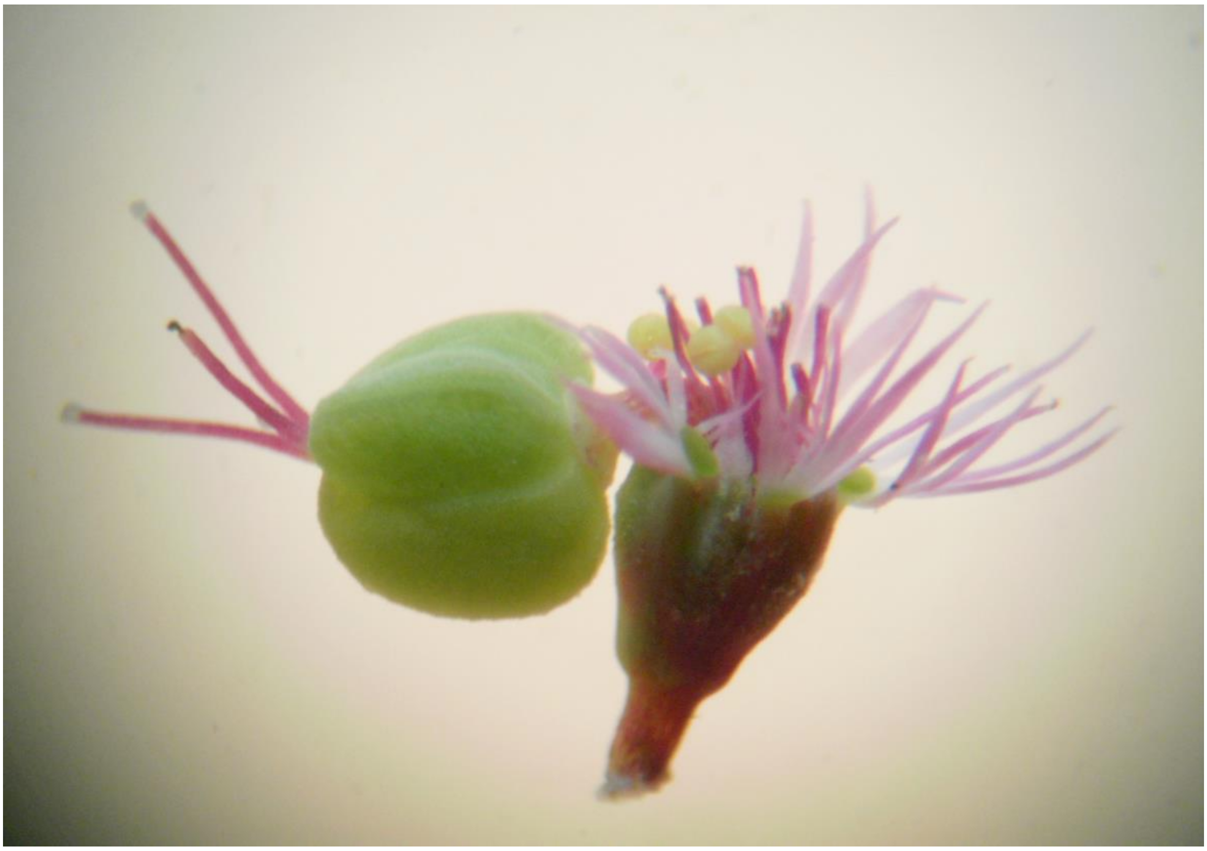

Mature cyathium

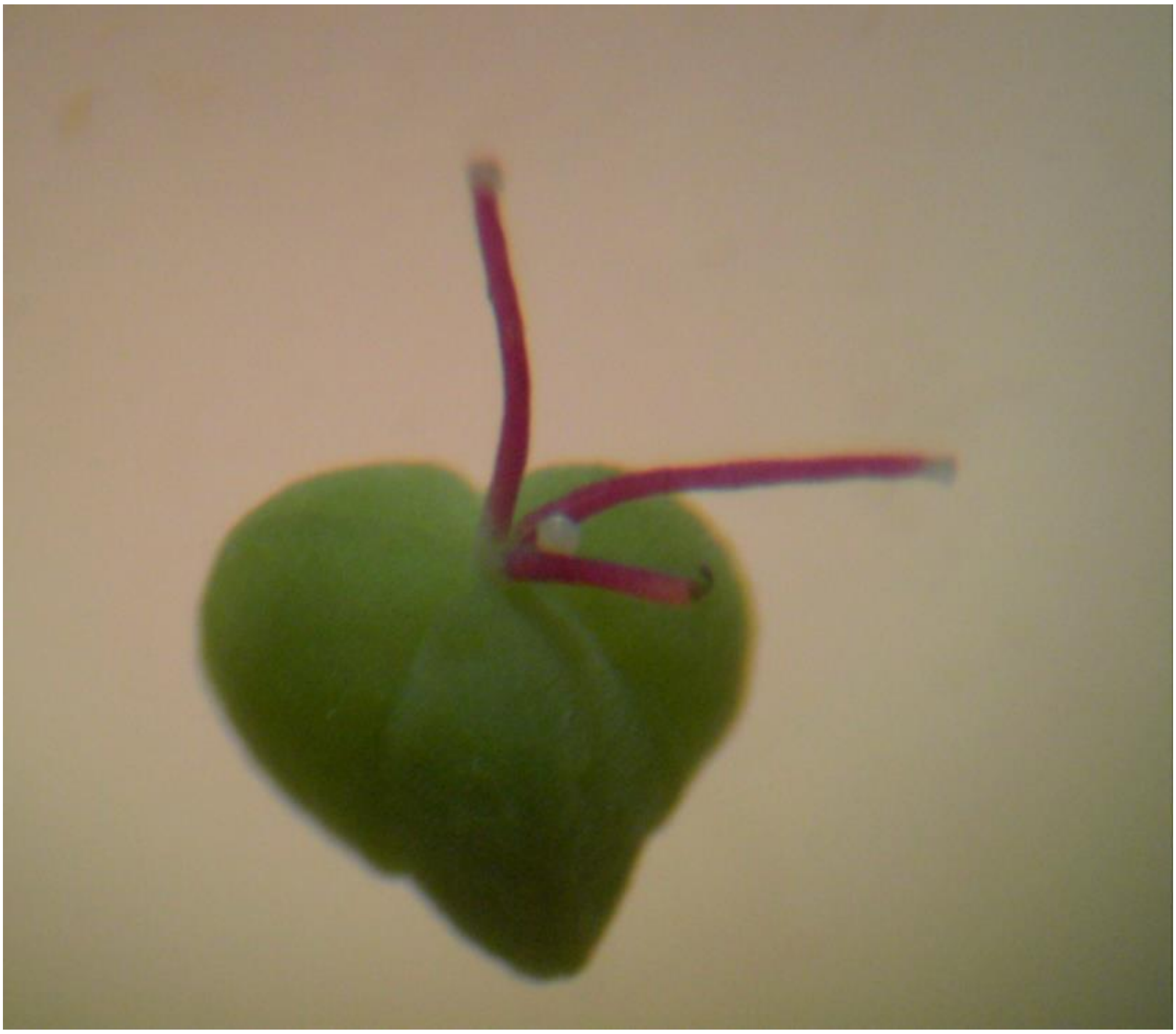

Capsule

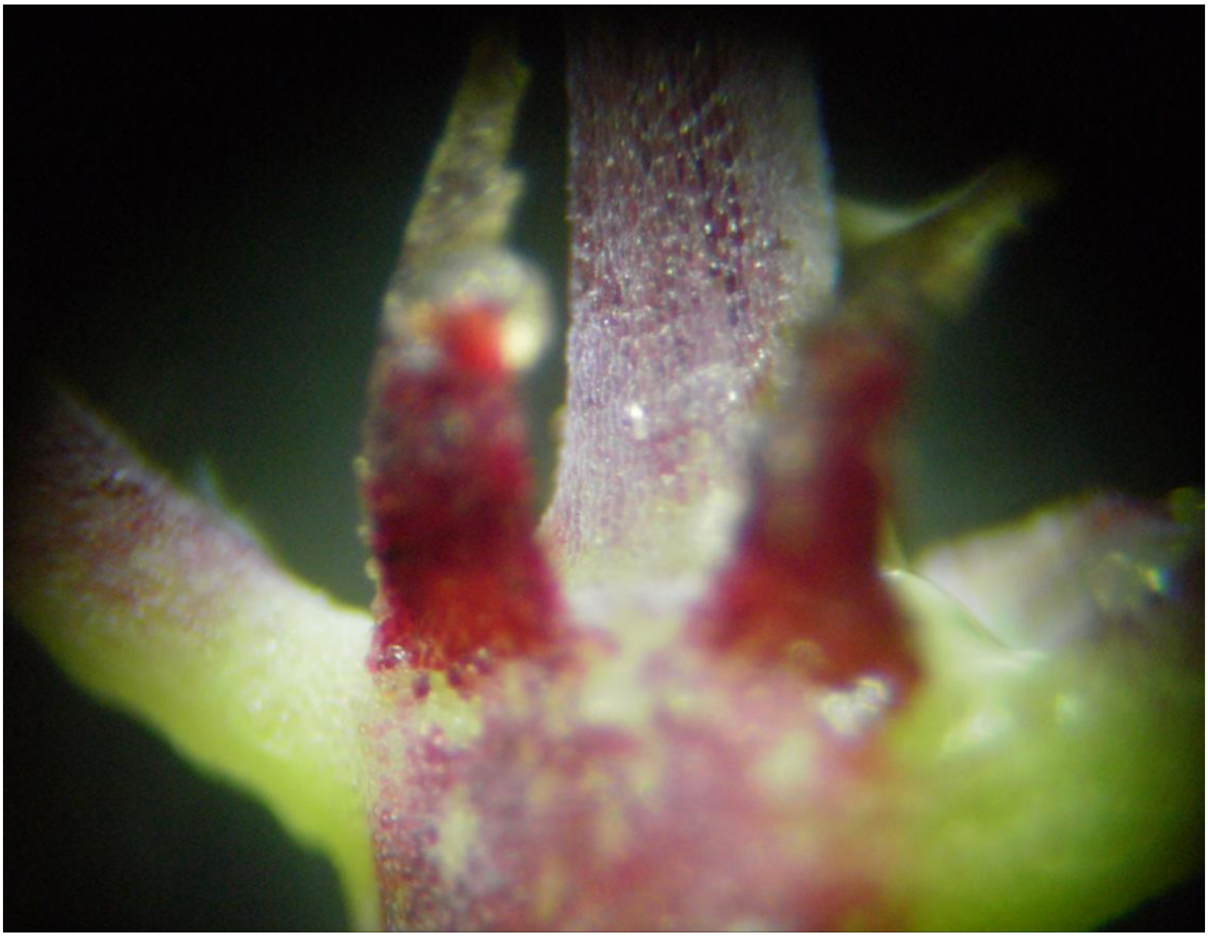

Stipules

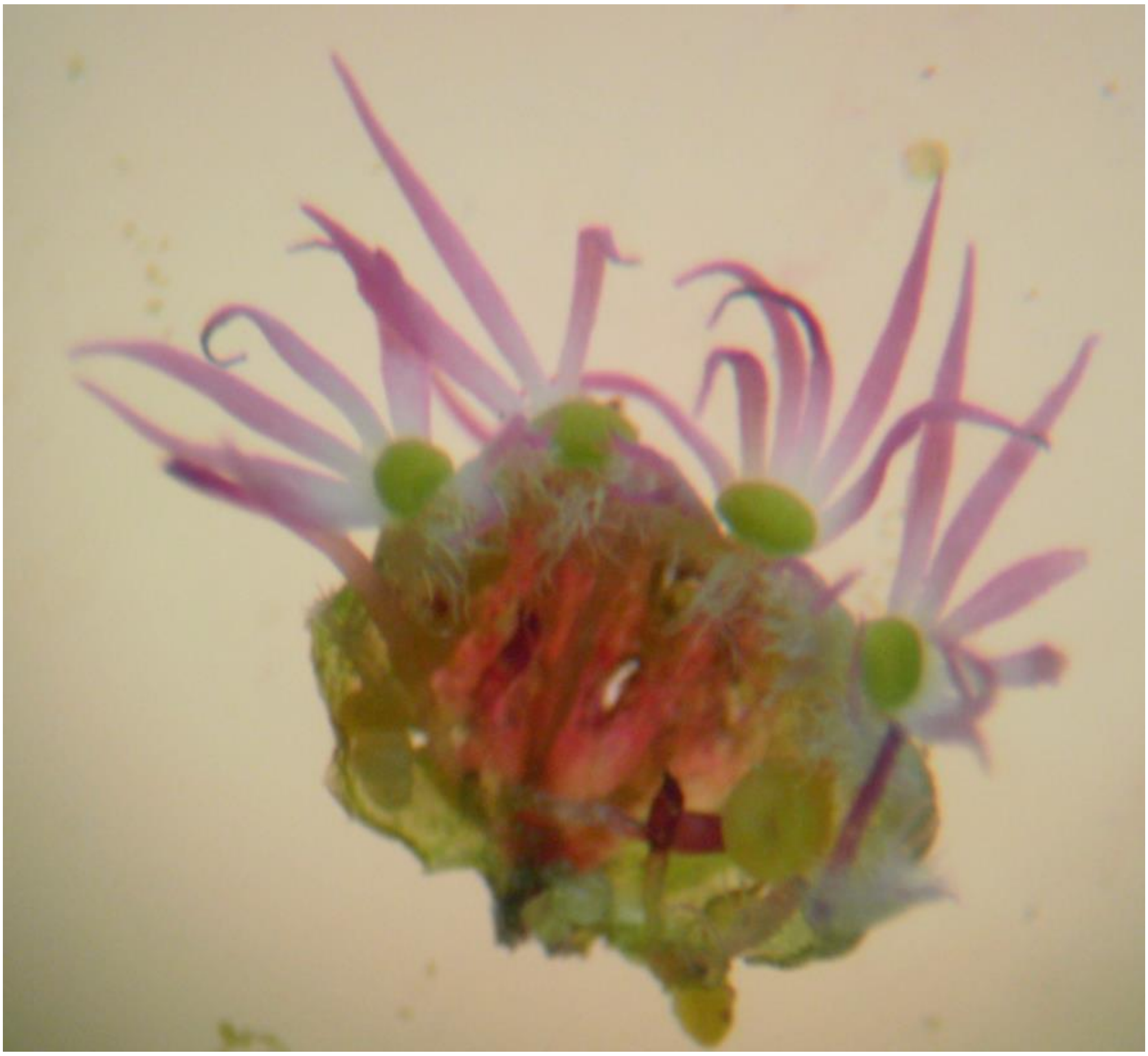

Portion of involucral cup showing lobes, glands and limbs
